# Embryonic type 3 innate lymphoid cells sense maternal dietary cholesterol to control local Peyer’s patch development

**DOI:** 10.1101/2023.03.19.533339

**Authors:** Kelsey Howley, Alyssa Berthelette, Simona Ceglia, Joonsoo Kang, Andrea Reboldi

**Affiliations:** Department of Pathology, University of Massachusetts Chan Medical School, Worcester, MA

## Abstract

Lymphoid tissue inducer (LTi) cells develop during intrauterine life and rely on developmental programs to initiate the organogenesis of secondary lymphoid organs (SLOs). This evolutionary conserved process endows the fetus with the ability to orchestrate the immune response after birth and to react to the triggers present in the environment. While it is established that LTi function can be shaped by maternal-derived cues and is critical to prepare the neonate with a functional scaffold to mount immune response, the cellular mechanisms that control anatomically distinct SLO organogenesis remain unclear.

We discovered that LTi cells forming Peyer’s patches, gut-specific SLOs, require the coordinated action of two migratory G protein coupled receptors (GPCR) GPR183 and CCR6. These two GPCRs are uniformly expressed on LTi cells across SLOs, but their deficiency specifically impacts Peyer’s patch formation, even when restricted to fetal window. The unique CCR6 ligand is CCL20, while the ligand for GPR183 is the cholesterol metabolite 7α,25-Dihydroxycholesterol (7α,25-HC), whose production is controlled by the enzyme cholesterol 25-hydroxylase (CH25H). We identified a fetal stromal cell subset that expresses CH25H and attracts LTi cells in the nascent Peyer’s patch anlagen. GPR183 ligand concentration can be modulated by the cholesterol content in the maternal diet and impacts LTi cell maturation in vitro and in vivo, highlighting a link between maternal nutrients and intestinal SLO organogenesis.

Our findings revealed that in the fetal intestine, cholesterol metabolite sensing by GPR183 in LTi cells for Peyer’s patch formation is dominant in the duodenum, the site of cholesterol absorption in the adult. This anatomic requirement suggests that embryonic, long-lived non-hematopoietic cells might exploit adult metabolic functions to ensure highly specialized SLO development in utero.

## Introduction

Peyer’s patches, secondary lymphoid organs (SLOs) on the antimesenteric side of the small intestine, ensure a 3D scaffold that orchestrates most of the adaptive immune processes in the gut, including the induction of immunoglobulin A^1–4^. Similar to other SLOs, the development of Peyer’s patch (PPs) is largely programmed during embryogenesis to allow efficient immune organization after birth^5–8^ and relies on lymphoid tissue inducer (LTi) cells to initiate SLO anlagen development. LTi cells belongs to the type 3 innate lymphoid cells (ILC3): despite sharing several transcription factors and surface receptors, LTi cells and ILC3 are generated from distinct progenitors in fetal liver^9,10^. In addition to LTi cells, PP induction requires lymphoid tissue initiator (LTin) cells, hematopoietic cells which express the receptor tyrosine kinase RET^11^. LTin cells signal to lymphoid tissue organizer (LTo) stromal cells to trigger chemokine production (CXCL13, CCL19, CCL21) and adhesion molecule expression (VCAM1, ICAM1, MADCAM1) that in turn attract LTi cells^12,13^. The resulting LTo-LTi interaction through LTbR-LTα1β2 results in increased levels of chemokines and adhesion molecules by LTo, driving follicle formation^14^.

LTi cells are required for all the SLOs, as demonstrate by the lack of lymph nodes (LNs) and PPs in mice deficient for the transcription factor RORgt and ID2^15–17^. However, differences between PP and LN organogenesis requirements are known: concomitant removal of T-bet and RORgt restores all SLO, but not PP, developement^18^, and a specific mutant of RORgt only impacts PPs^19^. Moreover, the developmental requirement for TRANCE and IL-7 signaling^20,21^ also suggests that a non-overlapping molecular program differentially controls PP and LN organogenesis.

The lymphoid organs development is considered to be imprinted during embryogenesis via distinct transcriptional programs, with LTin, LTi and LTo cells appearing in waves and giving rise to SLOs depending by cell-intrinsic ability to express chemokine and adhesion molecules. Nevertheless, data exist for the impact of the maternal diet on the secondary organ development^22^: these findings raise the possibility that environmental cues might shape lymphoid organogenesis through a cell-autonomous in utero signaling. However, so far, cell-extrinsic requirements for PP and LN organogenesis are limited to the widely studied retinoic acid signaling, and no additional dietary metabolite signaling has been shown to specifically control PP development.

Here we show that a cholesterol metabolite (i.e. oxysterol), 7α,25-HC, in concert with the chemokine CCL20, controls gut LTi to shape PP development in the duodenum. 7α,25-HC is the ligand for GPR183^23^, while CCL20 is the unique ligand for CCR6: both CCR6 and GPR183 are G-couple receptor proteins (GPCR) which mediate cell migration through Gai coupling. Despite being described on adult ILC3, GPR183 and CCR6 expression and role in LTi biology has not yet been elucidated.

Moreover, we found that cholesterol content of the maternal diet, but not maternal microbiome, impacts LTi maturation and PP development. Since intestinal cholesterol absorption is absent in the fetus, and the maternal-derived cholesterol reaches the fetal circulation through placental transport, our data suggest that local oxysterol production might differ in distinct intestinal areas. Therefore, discrete PPs along the small intestine might require locally distinct signals for their developments, highlighting the existence of an unreported, highly specific anatomical control of PP organogenesis.

### CCR6 and GPR183 are co-expressed by LTi cells and control LTi cell fate in Peyer’s patch anlagen

Lymphoid structures in the gut can be divided based on their developmental timing, cell requirements and positioning. PP development begins in the sterile uterine environment at E.14.5 and requires LTi cells: PPs are only present in the small intestine. In contrast, crypto patches (CP) and isolated lymphoid organs (ILF) development begins after birth and requires ILC3 cells and microbial stimuli: in contrast to PPs, CP and ILF are predominantly found in distal ileum and colon. GPR183 and CCR6 expression has been reported on ILC3 cells^24–26^ and single receptor deficiency leads to impaired CP and ILF development^25,27,28^. Interesting, no defect in PP development has been reported with either GPR183^-/-^ or CCR6^-/-^ mice, suggesting that fetal LTi cells might not express or require these receptors for their function.

We first analyzed the pattern of expression of CCR6 and GPR183 on fetal LTi cells using GPR183 KI/KO reporter mice^29^ in both small intestine and fetal liver. At E16.5, we could easily identify 3 distinct LTi cell populations based on CD4 and GPR183 expression in the gut (**Figure 1a,c**). CD4+ *Gpr183+* LTi cells were also uniformly high for IL7 receptor alpha (IL7Rα) and the mucosal integrin α4β7; CD4-*Gpr183+* LTi cells expressed intermediate level of IL7Rα and high level of α4β7 but showed bimodal level of CCR6 suggesting they might represent immature LTi cells^22^. CD4- *Gpr183*- LTi cells had low level of CCR6, but expressed intermediate level of IL7Rα and α4β7. Similar findings were observed in fetal liver, although the overall CD4+ *Gpr183+* population was reduced compared to small intestine (**Figure 1b**) possibly pointing to an expansion and/or maturation of the LTi subsets in the gut. Co-expression of *Gpr183* and CCR6 in mature LTi cells was not restricted to the uterine environment as day 1 (D1) LTi cells showed a similar pattern to the one observed in fetal liver and small intestine (**Suppl. Fig.1a,b,c**). It has been proposed that CD4-LTi cells might receive cues to differentiate into CD4+ LTi cells during secondary lymphoid organs development^22^: our data show that both CD4- and CD4+ LTi cells express *Gpr183*, in contrast CCR6 is highest in CD4+ LTi cells, suggesting that, while GPR183 precedes CCR6 expression, GPR183 and CCR6 co-expression marks mature LTi cells in neonatal PPs.

**Figure 1.**
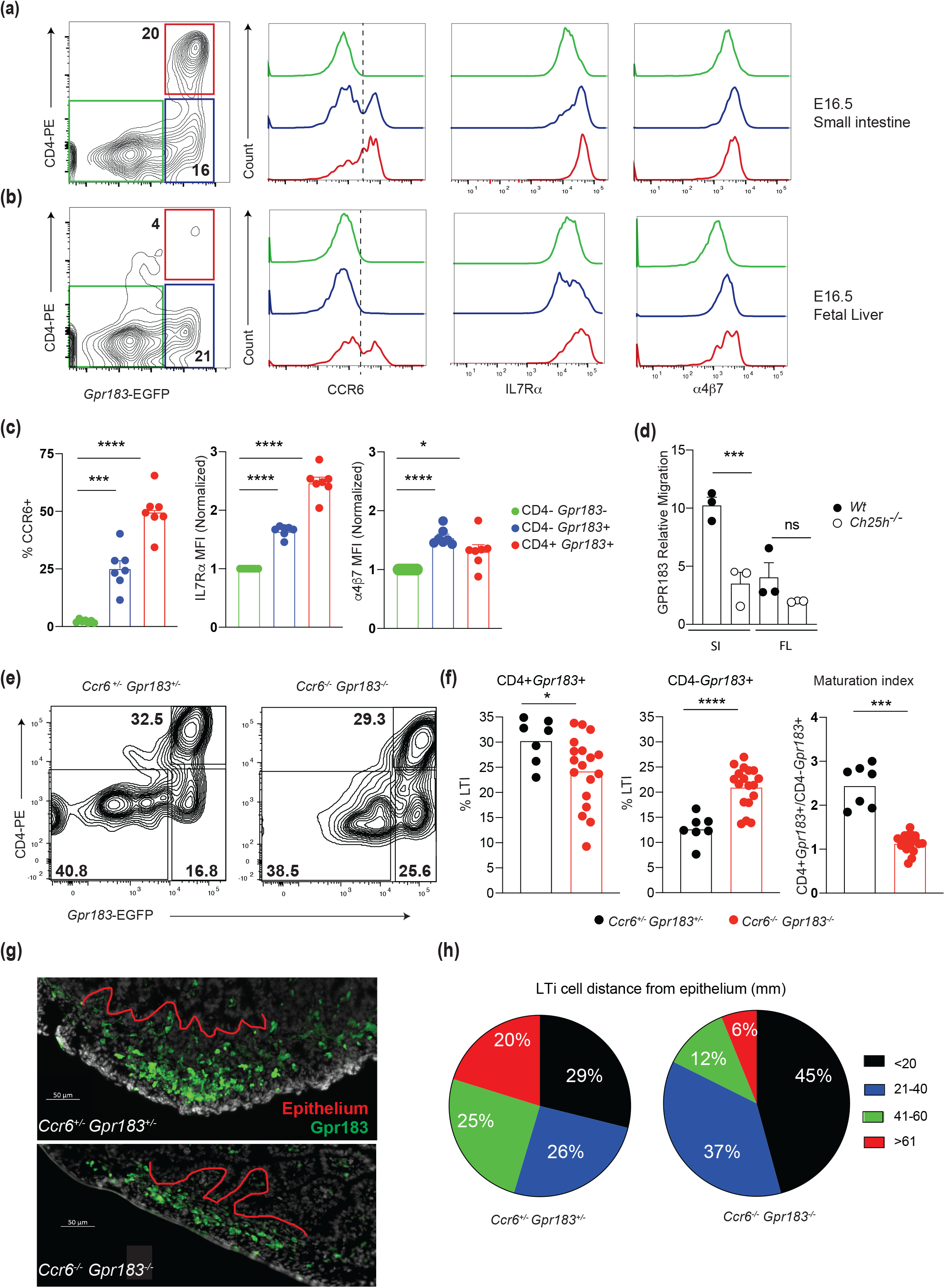
CCR6 and GPR183 are co-expressed by fetal LTi and required for their positioning and maturation. **a)** Representative flow cytometric contour plots and histograms depicting expression of *Gpr183*, CD4, CCR6, IL7Rα and α4β7 on CD45+ Lin- CD90+ cells in E15.5 small intestine from *Gpr183* reporter mouse. **b)** Representative flow cytometric contour plots and histograms depicting expression of Gpr183, CD4, CCR6, IL7Rα and α4β7 on CD45+ Lin- CD90+ cells in E15.5 liver from *Gpr183* reporter mouse. **c)** Compiled CCR6 frequency and IL7Rα and α4β7 MFI (normalized on CD4-*Gpr183*-) in different E15.5 small intestine LTi subsets gated as in a. **d)** Relative migration efficiency of GPR183 + cells to fetal small intestine and liver extracts. Tissue extract from *Ch25h^-/-^* were used as negative control. **e)** Representative flow cytometric contour plots and **e**) compiled data for the expression of *Gpr183* and CD4 on CD45+ Lin- CD90+ cells in E15.5 small intestine from *Ccr6^+/-^ Gpr183^+/-^* and *Ccr6^-/-^ Gpr183^-/-^*. **g)** Immunofluorescence analysis of E.15.5 small intestine from *Ccr6^+/-^ Gpr183^+/-^* and *Ccr6^-/-^ Gpr183^-/-^*. **h)** Pie charts showing the frequencies of LTi cells at different distance from the intestinal epithelium in *Ccr6^+/-^ Gpr183^+/-^* and *Ccr6^-/-^ Gpr183^-/-^*. Data are representative of at least three independent experiments. * p<0.05, ***p<0.005, **** p<0.001 as determined by as determined by Anova with Bonferroni correction (c) or unpaired t test (d,f).

To test weather fetal small intestine and liver contain GPR183 ligand (i.e. 7α,25-HC) we employed an in vitro bioassay that relies on GPR183-dependent cell line migration (**Suppl. Fig.1d**)^30–34^ and that has low nanomolar sensitivity for 7α,25-HC, but not for its precursor 25-HC (**Suppl. Fig.1e**). At E16.5 fetal small intestine extract drove GPR183 specific migration and therefore contained GPR183 ligand (**Figure 1d**). Cell migration in response to fetal liver tissue extract migration was indistinguishable from background migration using tissue extract from mice deficient for the enzyme CH25H and therefore unable to generate GPR183 ligand (**Figure 1d**).

Co-expression of migratory GPCR is often linked to in vivo cooperative migration to reach functional niche in primary and secondary lymphoid organs^30,34^, so we reasoned that CCR6 and GPR183 on LTi cells might work in concert to mediate LTi function. Thus, to assess the impact of both CCR6 and GPR183 on intestinal LTi cells we generated mice lacking both CCR6 and GPR183 (*Ccr6^-/-^ Gpr183^-/-^*, dKO). At E15.5 dKO small intestine showed reduced CD4+ *Gpr183+* LTi cells with expansion of CD4-*Gpr183+LTi* (Figure **1e**) and an overall impaired maturation (**Figure 1f**) compared to compound heterozygous mice (*Ccr6^+/-^ Gpr183^+/-^*, dHET). The overall frequency of LTi cells was not different between dHET and dKO, suggesting that these GPCRs, while contributing to local LTi maturation, do not mediate migration from the circulation into the intestine (**Sup. Fig.1f**). Interestingly, the relative contribution of mature and immature LTi cells was not maintained during the entire embryonic time, possibly suggesting discrete window for GPR183 and CCR6 function on LTi cells (**Sup. Fig.1g**). No difference between dHet and dKO was observed in fetal liver LTi cells. highlighting how these migratory receptors might be preferentially involved in intestinal control of LTi fate (**Sup. Fig.1g**). One caveat of flow cytometry analysis of fetal small intestine is that it does not allow for anatomical characterization of LTi cells involved in the lymphoid organogenesis of PPs. Therefore, we decided to visualize the PP analgen by immunofluorescent microscopy. While we could identify GPR183+ cells in the fetal anlagen of mice of both genotypes, mice lacking both CCR6 and GPR183 showed flat LTi cell cluster (**Figure 1g**), as measured by the distance from the intestinal epithelial layer (**Figure 1h**). Our data suggest that while LTi cells can migrate from the circulation to the intestinal wall in absence of both CCR6 and GPR183, they require these receptors to properly position in the nascent PP structure and undergo maturation.

### Anatomical zonation of Peyer’s patch development requires both CCR6 and GPR183

Altered fetal anlagen formation usually leads to defective SLO development. We reasoned that analysis of PPs in adult mice could provide insights on the defect observed in the embryonic gut with increased granularity. Therefore, we evaluatde adult co-house dHet and dKO for LN and PP structures. Peripheral LNs as well as mesenteric LNs were indistinguishable between dHet and dKO, however there was a significant reduction in PP number in mice lacking both CCR6 and GPR183 (**Supp Fig 2a**). Whole mounted small intestine^35^ revealed a very apparent altered distribution of PPs in dKO animals (**Figure 2a**): moreover, while PPs in dHET mice were characterized by multiple follicles, the remaining PPs in dKO mice tended to have a single, smaller follicle.

**Figure 2.**
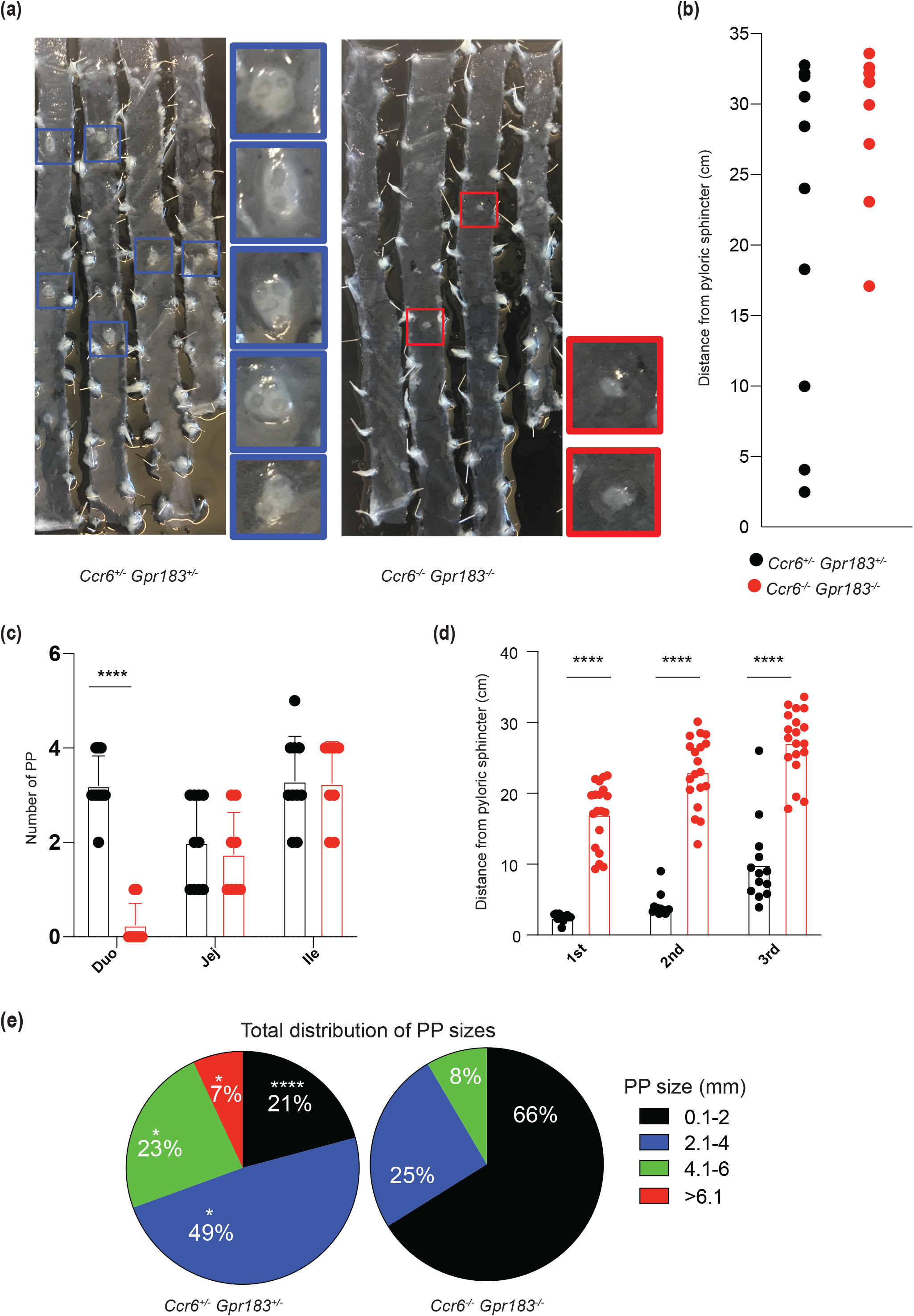
CCR6 and GPR183 deficiency abrogates PP development in the duodenum. **a)** Representative whole mount analysis of the small intestine from adult *Ccr6^+/-^ Gpr183^+/-^* and *Ccr6^-/-^ Gpr183^-/-^*. **b)** Representative distance from pyloric sphincter of Peyer’s patches (PPs) in adult *Ccr6^+/-^ Gpr183^+/-^* and *Ccr6^-/-^ Gpr183^-/-^*. **c)** Compiled number of PPs in different segment of the small intestine in adult *Ccr6^+/-^ Gpr183^+/-^* and *Ccr6^-/-^ Gpr183^-/-^*. **d)** Compiled distance from pyloric sphincter of the first three Peyer’s patches (PPs) in adult *Ccr6^+/-^ Gpr183^+/-^* and *Ccr6^-/-^ Gpr183^-/-^*. **e)** Pie charts showing the frequencies of PPs of different sizes in the small intestine from adult *Ccr6^+/-^ Gpr183^+/-^* and *Ccr6^-/-^ Gpr183^-/-^*. Data are representative of at least three independent experiments. * p<0.05, **** p<0.001 as determined by unpaired t test.

Lack of PPs was complete in the duodenal portion (**Figure 2b**), but with normal frequency in jejunum and ileum (**Figure 2c)**. DKO mice also developed the most proximal PPs well into the jejunum **2d**) and had an overall reduced PP size (**Figure 2e**). We occasionally found a structure in the duodenum of dKO mice (**Supp Fig 2b**), however the structure lacked classic B-cell T-cell zone division (**Supp Fig 2c**) and further flow cytometry analysis revealed a differently stark phenotype of immune cell subset compared to similarly located PP structure in dHet mice, with an increased frequency of activated lymphocytes (**Supp Fig 2d**). In order to assess whether CCR6 or GPR183 plays a dominant role in duodenal PP positioning, we analyzed PPs in mice lacking 3 of the alleles of interest (*Ccr6^-/-^ Gpr183^+/-^* and *Ccr6^+/-^ Gpr183^-/-^*). Despite subtle changes in location (**Supp Fig 2e**), neither of these animals phenocopied dKO mice in term of PP number, positioning of duodenal PPs or PP size (**Supp Fig 2f-h**). Together, our data suggest that both CCR6 and GPR183 are required to establish duodenal PP development.

### Uterine and LTi-restricted requirements for CCR6 and GPR183 function in Peyer’s patch development

During embryonic development, LTi cells represent the main population expressing CCR6 and GPR183, but in adult animals these receptors can be expressed by other hematopoietic cells. Most notably, B cells express CCR6 and GPR183^2,29^, albeit at different level depending by their activation status, and impaired B cells migration can lead to altered PPs. To assess whether the requirement for CCR6 and GPR183 was restricted during embryonic development for the formation of duodenal PPs, we took advantage of a combination of blocking reagents and conditional deficient mice. First, we used CCL20-Fc fusion proteins, which competes with the endogenous CCL20 for its binding to CCR6 and therefore blocks CCR6 function. We plugged a *Ccr6^-/-^ Gpr183^-/-^* with a female *Gpr183^+/-^*, so to generate pups with identical CCR6 level (*Ccr6^+/-^*) and variable GPR183 level (*Gpr183^-/-^* or *Gpr183^+/-^*). Pregnant dams were then injected with either CCL20-Fc or IgG-Fc once a day for three times starting at E14.5 (**Figure 3a**) and pups were analyzed at 6 weeks old for PP positioning, number and size. Temporal inhibition of CCR6 pathway in-utero coupled with GPR183 deficiency altered the PP positioning (**Figure 3b**) and placed the most proximal PP in the jejunum (**Figure 3c**). While PP numbers were also decreased (**Figure 3d**), PP size was not different between genotypes (**Figure 3e**), suggesting that CCR6 might play a role after birth to regulate PP size^36,37^. In contrast, mice treated with control IgG-Fc showed no difference in PP position or number (**Supp Fig 3a-e**). To further restrict GPR183 during this process, we generated mice lacking GPR183 in cell expressing RORgt (*Rorc^Cre^Gpr183^flox/flox^*)^38^, making them a de facto LTi conditional knock-out for GPR183 during uterine life. Pregnant dams were injected with CCL20-Fc or IgG-Fc as described above and the offspring were similarly analyzed at 6 weeks (**Figure 3f**). Simultaneous inability to sense CCR6 and GPR183 ligand by LTi cells in utero led to altered PP positioning (**Figure 3g**). PP number was decreased in the duodenum (**Figure 3h**), with the most proximal PP appearing towards the jejunum (**Figure 3i**). Similar to what was observed in GPR183^-/-^ animals that received CCL20-Fc in utero, PP size was largely unaffected (**Figure 3j**). Mice with the identical genotype treated with control IgG-Fc had no difference in PP position or number (**Supp Fig 3f-j**). Our data show that transient CCR6 inhibition coupled with LTi-specific GPR183 deficiency during embryonic development phenocopies what observed in mice globally lacking CCR6 and GPR183, suggesting that a developmental window exists for LTi expressing CCR6 and GPR183 to establish PP development in the duodenum.

**Figure 3.**
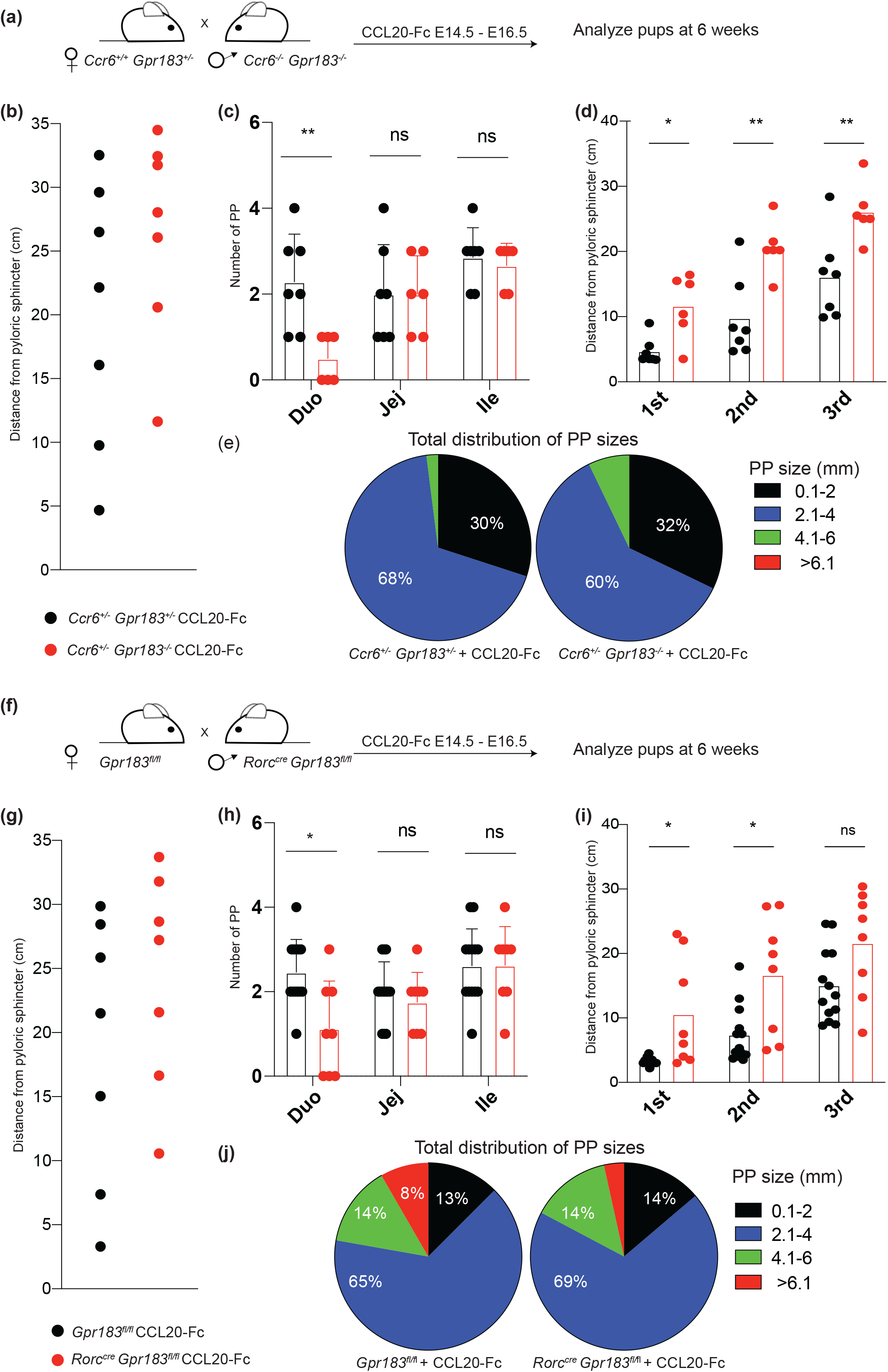
In utero alteration of CCR6 and GPR183 function impacts duodenal PPs. **a)** Schematic of the experimental approach to assess the effect of CCR6 inhibition in utero on PP development. **b)** Representative distance from pyloric sphincter of Peyer’s patches (PPs) in adult *Ccr6^+/-^ Gpr183^+/-^* and *Ccr6^+/-^ Gpr183^-/-^* treated in utero with CCL20-Fc. **c)** Compiled number of PPs in different segment of the small intestine in adult *Ccr6^+/-^ Gpr183^+/-^* and *Ccr6^+/-^ Gpr183^-/-^* treated in utero with CCL20-Fc. **d)** Compiled distance from pyloric sphincter of the first three Peyer’s patches (PPs) in adult *Ccr6^+/-^ Gpr183^+/-^* and *Ccr6^+/-^ Gpr183^-/-^* treated in utero with CCL20-Fc. **e)** Pie charts showing the frequencies of PPs of different sizes in the small intestine from adult *Ccr6^+/-^ Gpr183^+/-^* and *Ccr6^+/-^ Gpr183^-/-^* treated in utero with CCL20-Fc. **f)** Schematic of the experimental approach to assess the combined effect of CCR6 inhibition in utero and LTi-deficiency in GPR183 on PP development. **g)** Representative distance from pyloric sphincter of Peyer’s patches (PPs) in adult *Rorc^Cre^Gpr183^flox/flox^* and *Gpr183^flox/flox^* treated in utero with CCL20-Fc. **h)** Compiled number of PPs in different segment of the small intestine in adult *Rorc^Cre^Gpr183^flox/flox^* and *Gpr183^flox/flox^* treated in utero with CCL20-Fc. **i)** Compiled distance from pyloric sphincter of the first three Peyer’s patches (PPs) in adult *Rorc^Cre^Gpr183^flox/flox^* and *Gpr183^flox/flox^* treated in utero with CCL20-Fc. **j)** Pie charts showing the frequencies of PPs of different sizes in the small intestine from adult *Rorc^Cre^Gpr183^flox/flox^* and *Gpr183^flox/flox^* treated in utero with CCL20-Fc. Data are representative of at least three independent experiments. * p<0.05, ** p<0.01 as determined by unpaired t test.

### *Ch25h-expressing* cells provide essential GPR183 ligand in Peyer’s patch anlagen

The most potent ligand for GPR183 is 7α,25-HC which requires the enzyme CH25H as well as the enzyme CYP7B1^39^. To pinpoint whether GPR183 role in PP formation was centered around 7α,25-HC recognition, we generated mice which lacked both CCR6 and CH25H (*Ccr6^-/-^ Ch25h^-/-^*). DKO mice phenocopied mice lacking CCR6 and CH25H with specific reduction of duodenal PPs (**Figure 4a,b**). While single deficiency (*Ccr6^-/-^ Ch25h^+/-^* and *Ccr6^+/-^ Ch25h^-/-^*) slightly altered PP location in the small intestine (**Supp Fig 4a,b**), only dKO mice showed appearance of most proximal PP towards the jejunum (**Figure 4c, Supp Fig 4c**). Single deficient mice had normal PP number but slightly altered size (**Supp Fig 4d)** although not as profound as dKO (**Figure 4d)**. Our results proved that GPR183 in the fetal gut recognizes a metabolite which is generated by CH25H, i.e. 7α,25-HC, and, together with CCR6, drives PP development in the duodenum.

**Figure 4.**
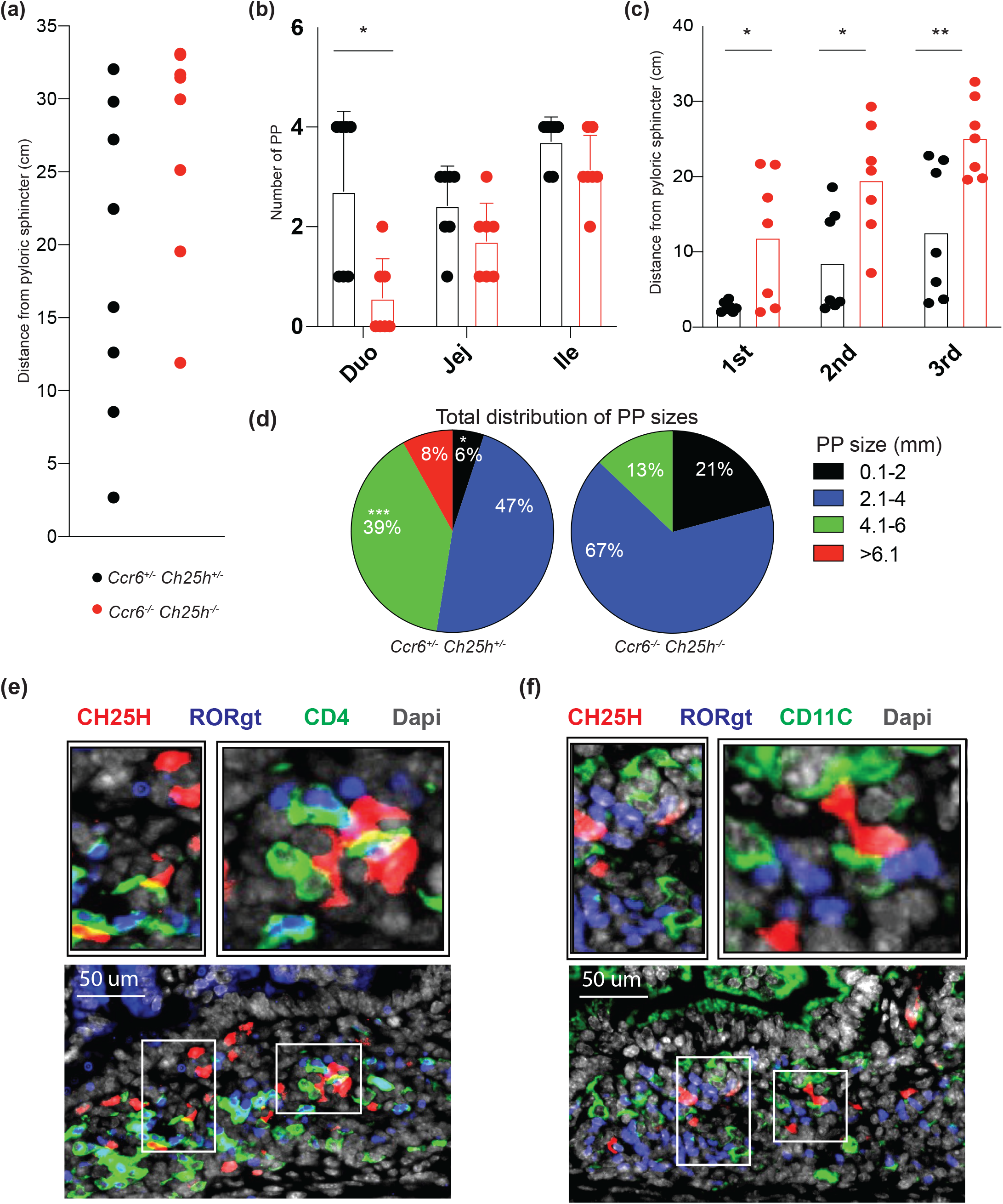
*Ch25h+* cells are required for GPR183 ligand generation and efficient PP development. **a)** Representative distance from pyloric sphincter of Peyer’s patches (PPs) in adult *Ccr6^+/-^ Ch25h^+/-^* and *Ccr6^-/-^ Ch25h^-/-^*. **b** Compiled number of PPs in different segment of the small intestine in adult *Ccr6^+/-^ Ch25h^+/-^* and *Ccr6^-/-^ Ch25h^-/-^*. **c)** Compiled distance from pyloric sphincter of the first three Peyer’s patches (PPs) in adult *Ccr6^+/-^* Ch25h^+/-^ and *Ccr6^-/-^ Ch25h^-/-^*. **d)** Pie charts showing the frequencies of PPs of different sizes in the small intestine from adult *Ccr6^+/-^ Ch25h^+/-^* and *Ccr6^-/-^ Ch25h^-/-^*. Immunofluorescence analysis of E.18.5 small intestine from *Ch25h*-reporter mouse stained with **g)** anti-RFP (red, capturing tdTom expression), anti-CD4, anti-RORgt antibodies, and DAPI; and **f)** anti-RFP, anti-CD11c and anti-RORgt antibodies, and DAPI. Data are representative of at least three independent experiments. * p<0.05, ** p<0.01, ***p<0.005 as determined by unpaired t test.

To visualize the GPR183 niche in vivo we took advantage of CH25H reporter mice^30^: analysis of E18.5 PP anlagen revealed the existence of *Ch25h+* cells, and they were found in close proximity to CD4+ RORγt+ cells, i.e. LTi cells (**Figure 4e)**. While CH25H can be expressed by myeloid cells^40^, in PP anlagen CH25H+ cells were negative for CD11c (**Figure 4f)** and instead co-expressed VCAM-1(**Supp Fig 4a,b**), a stromal cell marker that defines LTO^12^.

Together, our data revealed that cells with the enzymatic machinery necessary for the generation of the GPR183 ligand are present in the PP anlagen, and are required for the development of the duodenal PPs.

### TCF7 in LTi cells controls the generation of duodenal Peyer’s patches

Transcriptional control of LTi cells is critical to properly initiate SLO organogenesis, as observed in mice lacking LTi specific transcription factors (TFs)^16,18^, but these TFs are generally regarded as switches that imprint cell fate and are absolutely required for LTi presence at the site of all SLOs. However, our data suggest that migratory receptor activity can specifically underpins discrete anatomical zonation of PPs; therefore, we sought to establish whether this process relied on specific TFs. T cell factor 1 (TCF1), encoded by the gene *Tcf7*, is a key TF of early thymic progenitor differentiation^41,42^ and it is expressed at high levels by LTi precursors^9^. While *Tcf7^-/-^* mice have defects in various innate lymphocyte subsets^43,44^ but normal LN formation^45^, its role in PP organogenesis has not been fully characterized^43^.

We initially generated mice with TCF1 deficiency in hematopoietic cells by crossing *Tcf7^flox/flox^* to *Sox13^Cre^ (Sox13^Cre^ Tcf7^flox/flox^*) that deletes early during development in the hemogenic endothelium^46^. In these animals duodenal PPs were reduced in both number, position and size compared to littermate, co-housed controls (**Figure 5a-d)**, suggesting that TCF1 expression in hematopoietic cells controls local PP development. To restrict TCF1 deficiency to LTi cells, we crossed *Tcf7^flox/flox^* to *Rorc^Cre^* (*Rorc^Cre^Tcf7^flox/flox^*): we observed reduction in duodenal PPs number, altered position and smaller size compared to controls (**Figure 5e-h)**, a phenotype that fully recapitulates the one observed in mice lacking TCF1 in the whole hematopoietic compartment. Our findings suggest that TCF1 might be preferentially involved in the generation of duodenal PPs: whether this process requires TF activity at a precursor level in the fetal liver or directly in the small intestine remains to be determined.

**Figure 5.**
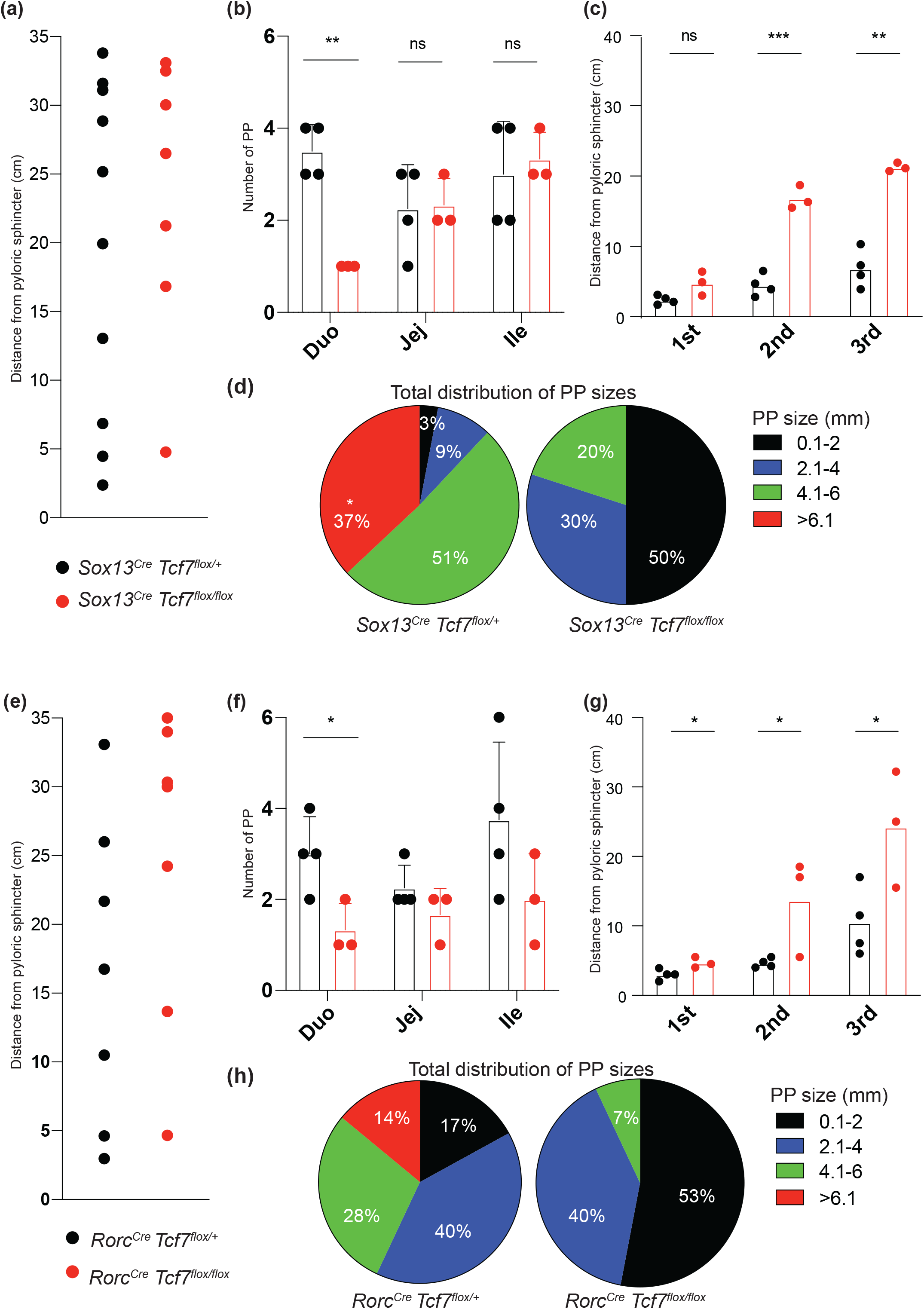
TCF1-expressing LTi cells control duodenal PP development. **a)** Representative distance from pyloric sphincter of Peyer’s patches (PPs) in adult *Sox13^Cre^Tcf7^flox/flox^* and *Sox13^Cre^Tcf7^flox/+^*. **b)** Compiled number of PPs in different segment of the small intestine in adult *Sox13_Cre_Tcf7^flox/flox^* and *Sox13^Cre^Tcf7^flox/+^*. **c)** Compiled distance from pyloric sphincter of the first three Peyer’s patches (PPs) in adult *Sox13^Cre^Tcf7^flox/flox^* and *Sox13^Cre^Tcf7^flox/+^*. **d)** Pie charts showing the frequencies of PPs of different sizes in the small intestine from adult *Sox13_Cre_Tcf7^flox/flox^* and *Sox13^Cre^Tcf7^flox/+^*. **e)** Representative distance from pyloric sphincter of Peyer’s patches (PPs) in adult *Rorc^Cre^Tcf7^flox/flox^* and *Rorc^Cre^Tcf7^flox/+^*. **f)** Compiled number of PPs in different segment of the small intestine in adult *Rorc^Cre^Tcf7^flox/flox^ and *Rorc^Cre^Tcf7^flox/+^*. **g)** Compiled distance from pyloric sphincter of the first three Peyer’s patches (PPs) in adult Rorc^Cre^Tcf7^flox/flox^* and *Rorc^Cre^Tcf7^flox/+^*. **h)** Pie charts showing the frequencies of PPs of different sizes in the small intestine from adult *Rorc^Cre^Tcf7^flox/flox^* and *Rorc^Cre^Tcf7^flox/+^*. Data are representative of at least three independent experiments. * p<0.05, ** p<0.01, ***p<0.005 as determined by unpaired t test.

### Differential impact of exogenous cues on LTi cell maturation and Peyer’s patch development

While the utero has long been considered a sterile environment, there is debate regarding whether commensals can be present, and whether they can have an impact on fetal immune development^47–50^.

To test if commensals modulation could affect PP organogenesis, we treated pregnant mother with single antibiotic (Ampicillin, Metronidazole or Vancomycin) during pregnancy starting at E.10.5 and discontinued them after birth. Antibiotic treatments had no impact on duodenal PP location or number (**Figure 6a-c**), although an effect was observed on overall size (**Figure 6d**), possibly due to altered post-birth commensal environment, despite dirty bedding from untreated mice being added to the cage post-partum.

**Figure 6.**
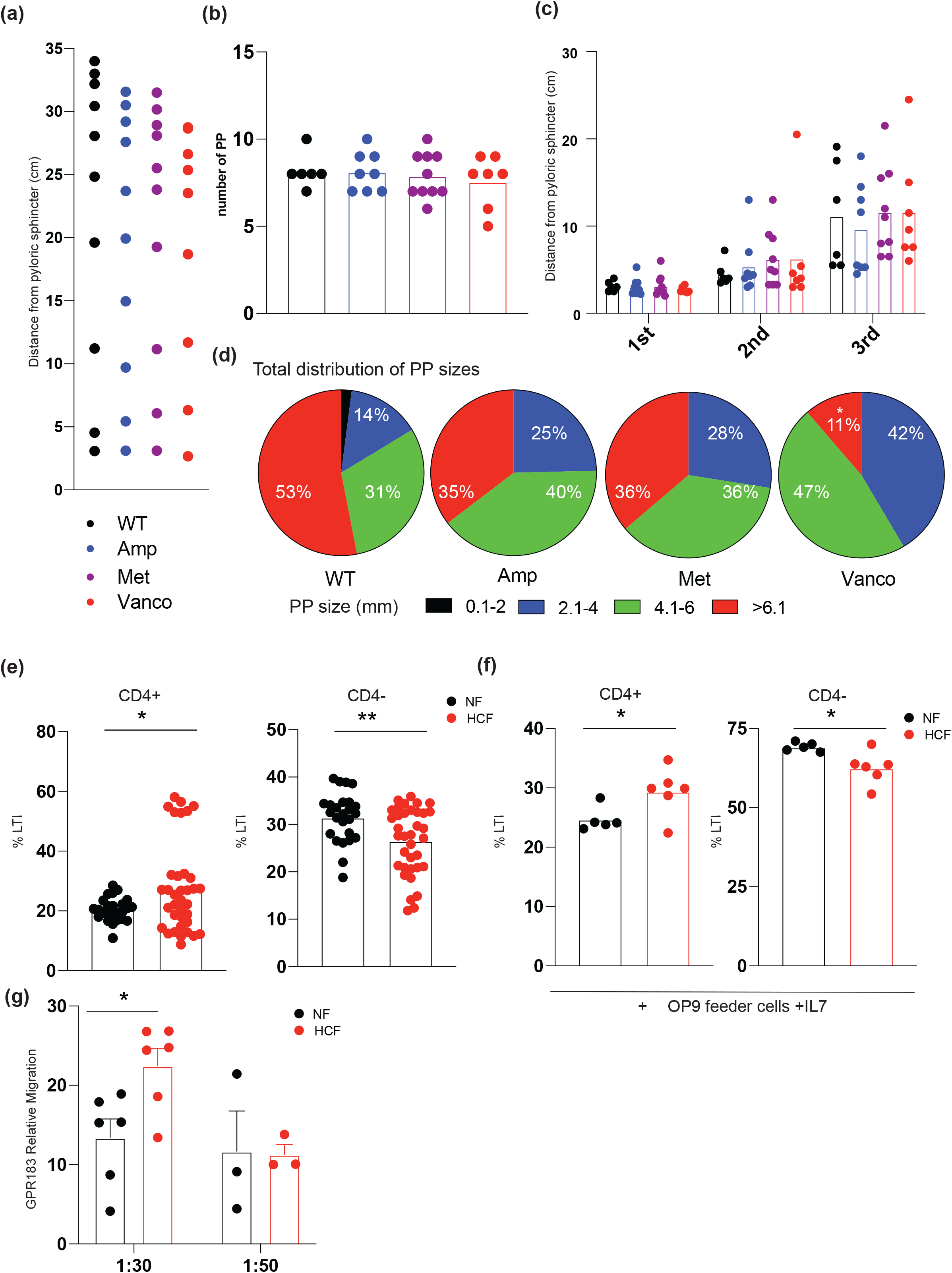
Dietary cholesterol, not commensal composition, impacts LTi cell maturation. **a)** Representative distance from pyloric sphincter of Peyer’s patches (PPs) in adult mice from dams treated with the single antibiotic Ampicillin, Metronidazole or Vancomycin only during pregnancy starting at E10.5. **b)** Compiled number of PPs in different segment of the small intestine in adult mice from dams treated with the single antibiotic Ampicillin, Metronidazole or Vancomycin only during pregnancy starting at E10.5. **c)** Compiled distance from pyloric sphincter of the first three Peyer’s patches (PPs) in adult mice from dams treated with the single antibiotic Ampicillin, Metronidazole or Vancomycin only during pregnancy starting at E10.5. **d)** Pie charts showing the frequencies of PPs of different sizes in the small intestine in adult mice from dams treated in utero with the single antibiotic Ampicillin, Metronidazole or Vancomycin only during pregnancy starting at E10.5. **e)** Compiled data for the frequency of CD4+ or CD4- CD45+ Lin- CD90+ cells in E16.5 small intestine of fetuses from dams fed either normal food (NF) or food with 2% cholesterol (HCF) starting at E.12.5 **f)** Compiled data for the frequency of CD4+ or CD4- LTi sorted from E16.5 small intestine of fetuses from dams fed either normal food (NF) or food with 2% cholesterol (HCF) starting at E.12.5 and culture for 3.5 days on OP9 feeder cells. **g)** Relative migration efficiency of GPR183 + cells to fetal small intestine extracts of fetuses from dam fed either normal food (NF) or food with 2% cholesterol (HCF). Data are representative of at least three independent experiments. * p<0.05, ** p<0.01 as determined by unpaired t test.

In utero, fetuses are exposed to dietary components of the maternal diet, so we tested whether increase in cholesterol-content in the maternal diet could affect LTi cells. We fed pregnant dam with food with the sole addition of 2% cholesterol, or control food, from E. 12.5 to E.16.5 and then analyzed the LTi population by flow cytometry. Fetuses which were exposed to cholesterol diet showed an increase in CD4+ LTi cell fraction (i.e. mature LTi cells) in the small intestine compared to fetuses from dam fed control food (**Figure 6e**). LTi cells sorted from E16.5 intestine of dams exposed to different food and cultured in vitro with OP9 and IL7, an efficient way to maintain ILC3 cells in vitro^51,52^, showed that LTi cell primed by high cholesterol maternal diet had increased propensity to assume and maintain mature status (**Figure 6f**). Analysis of tissue 7α,25-HC using in vitro bioassay confirmed that increased dietary cholesterol uptake from maternal side led to an enhanced production of GPR183 ligand (**Figure 6g**). Together, our findings showed that maternal diet can directly impact fetal immune cells by generating a cholesterol metabolite directly in the fetal gut. The intestinal immune system is especially sensitive to nutritional status: PP counts and intestinal IgA are altered in underfed infants^53,54^; and immune response to oral vaccines, but not to injectable vaccines, is rapidly blunted in children from malnourished mothers^55–57^. While these children show widespread immune defects^58,59^, one enticing possibility is that maternal malnutrition impacts intestinal neonate SLO formation. In line with this concept, at least one maternal-derived metabolite has shown to be involved in PP organogenesis^22^ and responses to oral challenges^60^.

Our data show that complete PP organogenesis requires cooperation between two distinct migratory G protein coupled receptors (GPCRs) CCR6 and GPR183. The ligand for GPR183 is the cholesterol metabolite 7α,25-HC: the enzyme required for its production is present in the PP anlagen and maternal nutrition can modulate this metabolite concentration in the fetal gut.

The fetal intestine does not absorb nutrients, so the fetus receives all the necessary nutrients through blood vessels in the umbilical cord. However, fetal intestinal cells will eventually be exposed to luminal nutrients once outside the womb, so we speculate that fetal cells in the PP anlagen anticipate adult tissue function by integrating maternal-derived environmental cues, a process that has been described for other immune cells in barrier tissues^30^. We recently showed that in adult animals, dietary cholesterol impact mucosal immunity via oxysterol production^31,32^: we hypothesize that immune response in the intestine is exquisitely sensitive to changes in nutrition status, even during development. Our data show that maternal dietary cholesterol reaches the fetal gut and is metabolized by a specific fetal stromal cell subset to produce immunomodulatory cholesterol byproducts, creating a migratory and metabolic depot that instructs LTi maturation and PP organogenesis initiation. Why this process is dominant in the duodenum, and whether distinct nutrient sensing pathways can imprint PP development in other section of the small intestine is not known, however, differences in immune cell function associated to discrete anatomical locales along the gut has been reported before^31,61–64^. Additional investigation will be needed to elucidate the detailed contribution of maternal diet components to the development of mucosal immune system, possibly to generate interventions geared toward enhanced immunity at the barrier tissues during early life^65^.

## ACKNOWLEDGMENTS

We thank Daniel Campbell for the CCL20-Fc plasmid, Lawrence Stern and Grant Weaver for the generation of CCL20-Fc and IgG-Fc, and Fiona Raso for critical reading of the manuscript.

This work was supported by a Kenneth Rainin Foundation Innovator Award grant and by National Institutes of Health (NIH) grants AI155727 and AI171474 to A.R. S.C. was supported by the American Association of Immunologists Careers in Immunology Fellowship Program and Charles A. King Trust Postdoctoral Research Fellowship Award.

## AUTHOR CONTRIBUTIONS

K.H and A.R. conceived the study, developed the concept, designed the experiments, and wrote the paper. K.H, A.B and S.C and A.R. performed the experiments, analyzed the data, and interpreted the results, J.K. interpreted the results. All the authors contributed to the review and editing of the manuscript. A.R. reviewed the data and supervised the research.

