## Supplementary information for "Embryonic type 3 innate lymphoid cells sense maternal dietary cholesterol to control local Peyer’s patch development"

### MATERIALS AND METHODS

**Mice.** Ccr6$^{-/-}$ (Stock no: 005793), *Ch25h*$^{-/-}$ (Stock no: 016263), C57BL/6J mice (Stock no: 000664), *Rorc*$^{Cre}$ mice (Stock no: 022791) were from Jax Laboratories. *Gpr183*$^{+/Egfp}$ (reporter insertion creates a null allele), *Gpr183*$^{fl/fl}$ , *Tcf7*$^{flox/flox}$, *Ch25h*$^{tdTom}$ mice were described previously[29,30,38,66].

Males and females were used for experiments, but sex-matched within an experiment. No differences were observed between sexes. Ages of mice used for experiments are indicated in Figure Legends. Animals were randomly allocated to experimental groups. All mouse procedures were approved by the University of Massachusetts Medical School IACUC

**Diets.** Mice were either fed a standard chow diet (Prolab IsoPro RMH 3000 5P76) or a high cholesterol diet (2% cholesterol added to the Prelab RMH 3000 5P76; Envigo TD. 200179, customized diet) for the duration of the experiment.

**Peyer's patches measurement.** Analysis of whole mount small intestine was performed as described[35]. In brief, small intestines were removed intact, flushed with ice-cold phosphate-buffered saline (PBS), and opened along the mesenteric border. Intestines were pin on wax plates lumen facing up, and incubated in Hank's balanced salt solution (HBSS) + 5 mM EDTA for 12 min at 37°C shaking (90 rpm). After a wash in ice-cold PBS, small intestines were placed in 10% formalin-buffered saline at 4°C for overnight. The tissue was washed in 1 M NaCl, 1 M Tris, pH 7.2, and 0.5% Triton X-100 (TBST) three times and incubated in 1% H2O2 in methanol for 15 min. PP position was measured from the pyloric sphincter with a ruler.

**Cell Isolation.** The small intestine was opened longitudinally, washed with complete medium (10% heat-inactivated FBS, 10 mM Hepes, pH 7.2 and 1% penicillin–streptomycin), chopped with scissors and digested at 37 °C for 20 min in digestion medium (RPMI medium, 5% heat-inactivated FBS, 10 mM Hepes, pH 7.2, 0.5 mg ml/1 of Collagenase IV (Worthington Biochemical, catalog no. LS004189) and 50 µg/ml of DNase I (Sigma-Aldrich, catalog no. DN25)) in agitation Digested tissue was passed through a 70-µm cell strainer and isolated cells were stained for flow cytometry analysis. Fetal liver was smashed passed through a 70-µm cell strainer, undergone RBC lysis and the isolated cells were stained for flow cytometry analysis.

**Antibodies and flow cytometry.** Flow cytometry staining was performed in 96-well microtiter plates. Antibody cocktails were diluted in PBS (GIBCO) + 2% FBS (Sigma-Aldrich) + 2 mM EDTA (Teknova), and cells were stained in 50 mL for 20 min on ice. All the antibodies were from Biolegend. All samples were labeled with Fixable Viability Dye (Thermofisher) to exclude deal cells from all analysis. Data were acquired on a BD LSRII cytometer or FACSAria (BD Biosciences) and analyzed using FlowJo (Treestar).

**Cell sorting.** For LTi cells, small intestine from E16.5 fetuses were stained and sorted as Live CD3- CD11c- B220- CD19- CD45+ CD90+ cells with a 70mm on a BD FACSAria II.

**Cell culture.** $1.5 \times 10^3$ sorted LTi cells were cultured in 96-well plate with $1.2 \times 10^4$ OP9 cells per well (American Type Culture Collection) and 10 ng/ml recombinant mouse IL-7 (R&D Systems) in RPMI 1640 medium (supplemented with HEPES, FCS, sodium pyruvate, 2-mercaptoethanol, streptomycin-penicillin and l-glutamine).

**Immunofluorescence.** For immunofluorescence, tissues were fixed in 4% paraformaldehyde (J.T. Baker),0.53 M L-Lysine, 2.1 mg/ml sodium m-periodate, 0.024 N NaOH in PB for at least 2 hours at 4C, washed three times for 10 min in PB, then moved to 30% sucrose in PBS overnight. Tissues were flash frozen in Tissue-Tek Cryomold (VWR) the next day, and 7-mm sections were cut and then dried for 1 hour before staining. Sections were rehydrated in PBS, blocked, and then stained in primary antibody overnight at 4 C and stained for subsequent steps for 2 hours at room temperature. Slides were blocked with 5% normal mouse serum + 5% normal rat serum in PBS, 0.3% Triton-X100 (Sigma), 0.2% Bovine Serum Albumin (BSA) and 0.1% sodium azide. All the antibodies were diluted in the same buffer containing 2% normal mouse serum + 2% normal rat serum. Images were obtained with a Zeiss AxioObserver.Z1 (Carl Zeiss) inverted microscope and were analyzed by imaging processing software ImageJ (NIH).

**Oxysterol quantification.** Oxysterols- Lipids from tissues and IECs were extracted using the Folch method for lipid extraction. Briefly, tissues were weighed, homogenized in serum-free medium containing 0.5% BSA and prepared at a concentration of 100 mg ml−1. The 7α,25-HC and the 25HC activity was evaluated by transwell chemotaxis assay. Thus, lipid extracts were diluted in 10 volumes of sterile chemotaxis medium (RPMI + 0.5% fatty acid-free BSA) and tested for GPR183-dependent bioactivity by seeding on transwell, 50:50 mixed, M12 B cell line transduced with an GPR183–IRES–GFP retroviral construct and mock M12 cells. The migration assay was performed at 37 °C for 3 h and cells were analyzed by flow cytometry. The migration of GPR183–GFP+ M12 cells over M12 cells (which indicate the relative concentration of the GPR183 ligand, 7α,25-HC) was normalized to the migration toward lipid-free migration medium. The purified 7α,25-HC was used as a positive control at a concentration of 100 nM.

**LEGENDS**

**Supplementary Figure 1**. **CCR6 and GPR183 are co-expressed by neonatal LTi.**

**a)** Representative flow cytometric contour plots and histograms depicting expression of *Gpr183*, CD4, CCR6, IL7Rα and α4β7 on CD45+ Lin- CD90+ cells in D1 small intestine from *Gpr183* reporter mouse. **b)** Representative flow cytometric contour plots and histograms depicting expression of Gpr183, CD4, CCR6, IL7Rα and α4β7 on CD45+ Lin- CD90+ cells in D1 liver from *Gpr183* reporter mouse. **c)** Compiled CCR6 frequency and IL7Rα and α4β7 MFI (normalized on CD4- *Gpr183*-) in different E15.5 fetal liver LTi subsets. **d)** Schematic for GPR183-mediated Transwell migration assay. **e)** Relative migration efficiency of GPR183 + cells to synthetic 7α,25-HC and 25-HC. **f)** Compiled data for the frequency of total CD90+cells (among CD45+ Lin- cells) and *Gpr183* and CD4 (among CD45+ Lin- CD90+ cells) in small intestine and fetal liver at different embryonic time point from *Ccr6*$^{+/-}$ *Gpr183*$^{+/-}$ and *Ccr6*$^{-/-}$ *Gpr183*$^{-/-}$.

Data are representative of at least three independent experiments. ** p<0.01, ***p<0.005, **** p<0.001 as determined by Anova with Bonferroni correction (c) or unpaired t test (e)

**Supplementary Figure 2. CCR6 and GPR183 combined deficiency does not globally reduce SLOs and single deficiency has no impact on PP development.**

**a)** Compiled analysis of SLOs from adult *Ccr6*$^{+/-}$ *Gpr183*$^{+/-}$ and *Ccr6*$^{-/-}$ *Gpr183*$^{-/-}$. **b)** Compiled frequency of adult *Ccr6*$^{+/-}$ *Gpr183*$^{+/-}$ and *Ccr6*$^{-/-}$ *Gpr183*$^{-/-}$. with first PP analgen at different distance from pyloric sphincter. **c)** Representative immunofluorescent of the first aggregate found in small intestine of adult *Ccr6*$^{+/-}$ *Gpr183*$^{+/-}$ and *Ccr6*$^{-/-}$ *Gpr183*$^{-/-}$ stained for IgD, CD3, Gpr183 and DAPI. **d)** Compiled flow cytometry data regarding immune cells isolated from the excised first aggregate found in small intestine of adult *Ccr6*$^{+/-}$ *Gpr183*$^{+/-}$ and *Ccr6*$^{-/-}$ *Gpr183*$^{-/-}$. **e)** Representative distance from pyloric sphincter of Peyer's patches (PPs) in adult *Ccr6*$^{-/-}$ *Gpr183*$^{+/-}$ and *Ccr6*$^{+/-}$ *Gpr183*$^{-/-}$. **f)** Compiled number of PPs in different segment of the small intestine in adult *Ccr6*$^{-/-}$ *Gpr183*$^{+/-}$ and *Ccr6*$^{+/-}$ *Gpr183*$^{-/-}$. **g)** Compiled distance from pyloric sphincter of the first three Peyer's patches (PPs) in adult *Ccr6*$^{-/-}$ *Gpr183*$^{+/-}$ and *Ccr6*$^{+/-}$ *Gpr183*$^{-/-}$. **h)** Pie charts showing the frequencies of PPs of different sizes in the small intestine from adult *Ccr6*$^{-/-}$ *Gpr183*$^{+/-}$ and *Ccr6*$^{+/-}$ *Gpr183*$^{-/-}$. Data are representative of at least three independent experiments. * p<0.05, ** p<0.01, ***p<0.005, **** p<0.001 as determined by unpaired t test.

**Supplementary Figure 3. In utero injection of Fc protein has no effect on PP development.**

**a)** Schematic of the experimental approach to control for Fc protein in utero. **b)** Representative distance from pyloric sphincter of Peyer's patches (PPs) in adult *Ccr6*$^{+/-}$ *Gpr183*$^{+/-}$ and *Ccr6*$^{+/-}$ *Gpr183*$^{-/-}$ treated in utero with IgG-Fc. **c)** Compiled number of PPs in different segment of the small intestine in adult *Ccr6*$^{+/-}$ *Gpr183*$^{+/-}$ and *Ccr6*$^{+/-}$ *Gpr183*$^{-/-}$ treated in utero with IgG-Fc. **d)** Compiled distance from pyloric sphincter of the first three Peyer's patches (PPs) in adult *Ccr6*$^{+/-}$ *Gpr183*$^{+/-}$ and *Ccr6*$^{+/-}$ *Gpr183*$^{-/-}$ treated in utero with IgG-Fc. **e)** Pie charts showing the frequencies of PPs of different sizes in the small intestine from adult *Ccr6*$^{+/-}$ *Gpr183*$^{+/-}$ and *Ccr6*$^{+/-}$ *Gpr183*$^{-/-}$ treated in utero with IgG-Fc. **f)** Schematic of the experimental approach to assess the combined effect of CCR6 inhibition in utero and LTi-deficiency in GPR183 on PP development. **g)** Representative distance from pyloric sphincter of Peyer's patches (PPs) in adult *Rorc*$^{Cre}$*Gpr183*$^{flox/flox}$ and *Gpr183*$^{flox/flox}$ treated in utero with IgG-Fc. **h)** Compiled number of PPs in different segment of the small intestine in adult *Rorc*$^{Cre}$*Gpr183*$^{flox/flox}$ and *Gpr183*$^{flox/flox}$ treated in utero with IgG-Fc. **i)** Compiled distance from pyloric sphincter of the first three Peyer's patches (PPs) in adult *Rorc*$^{Cre}$*Gpr183*$^{flox/flox}$ and *Gpr183*$^{flox/flox}$ treated in utero with CCL20-Fc. **j)** Pie charts showing the frequencies of PPs of different sizes in the small intestine from adult *Rorc*$^{Cre}$*Gpr183*$^{flox/flox}$ and *Gpr183*$^{flox/flox}$ treated in utero with IgG-Fc. Data in are representative of at least three independent experiments.

**Supplementary Figure 4**. ***Ch25h*+ cells in PP anlagen are VCAM+ and are required for GPR183 ligand generation and efficient PP development.**

**a)** Representative distance from pyloric sphincter of Peyer's patches (PPs) in adult *Ccr6*$^{-/-}$ *Ch25h*$^{+/-}$ and *Ccr6*$^{+/-}$ *Ch25h*$^{-/-}$. **b** Compiled number of PPs in different segment of the small intestine in adult *Ccr6*$^{-/-}$ *Ch25h*$^{+/-}$ and *Ccr6*$^{+/-}$ *Ch25h*$^{-/-}$. **c)** Compiled distance from pyloric sphincter of the first three Peyer's patches (PPs) in adult *Ccr6*$^{-/-}$ *Ch25h*$^{+/-}$ and *Ccr6*$^{+/-}$ *Ch25h*$^{-/-}$. **d)** Pie charts showing the frequencies of PPs of different sizes in the small intestine from adult *Ccr6*$^{-/-}$ *Ch25h*$^{+/-}$ and *Ccr6*$^{+/-}$ *Ch25h*$^{-/-}$. **e)** Immunofluorescence analysis of E.18.5 small intestine from double *Ch25h*-reporter and *Gpr183*-reporter mouse stained with anti-RFP (red, capturing tdTom expression), and anti-GFP and anti-VCAM1 antibodies, and DAPI.

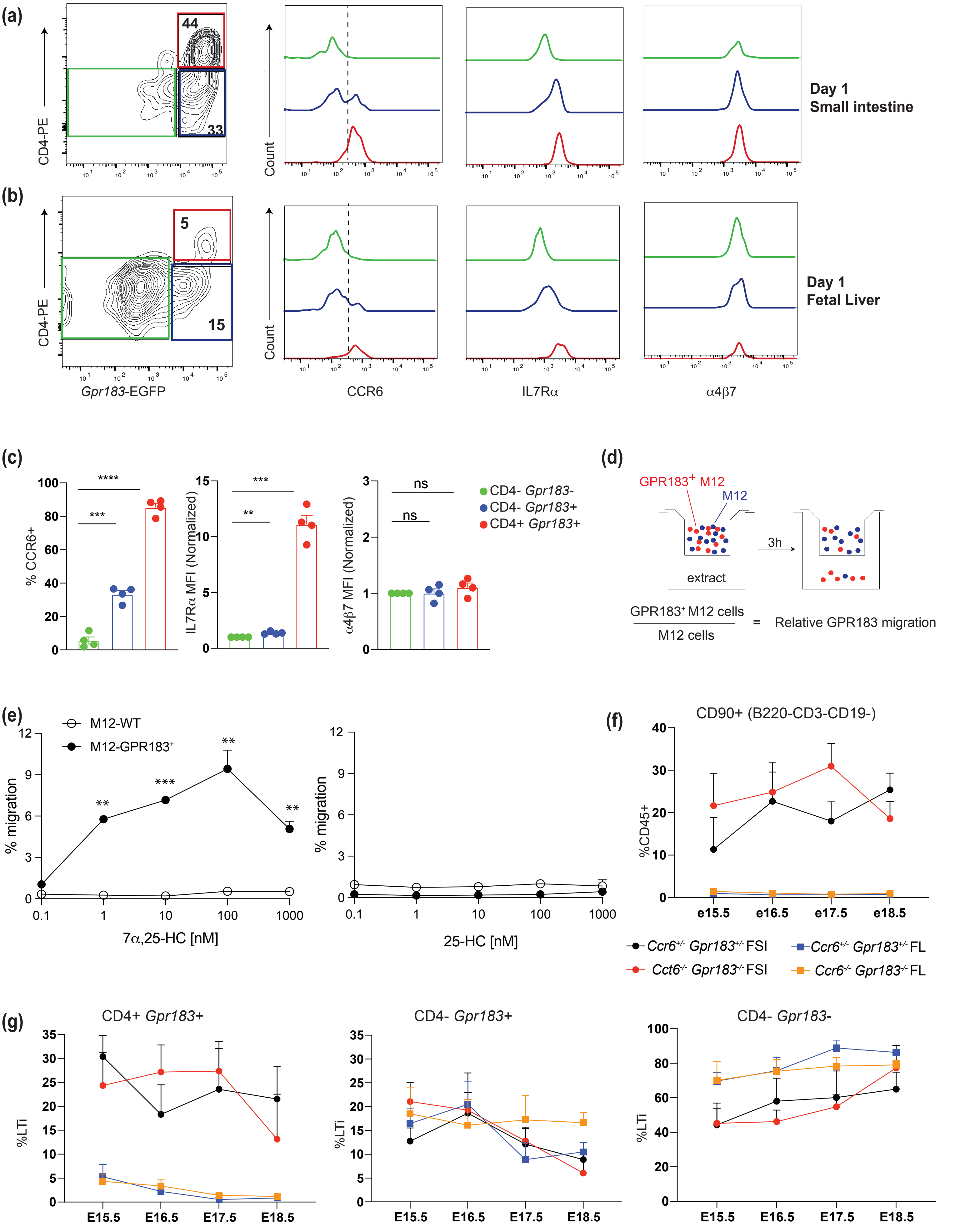


**Supplementary Figure 1**

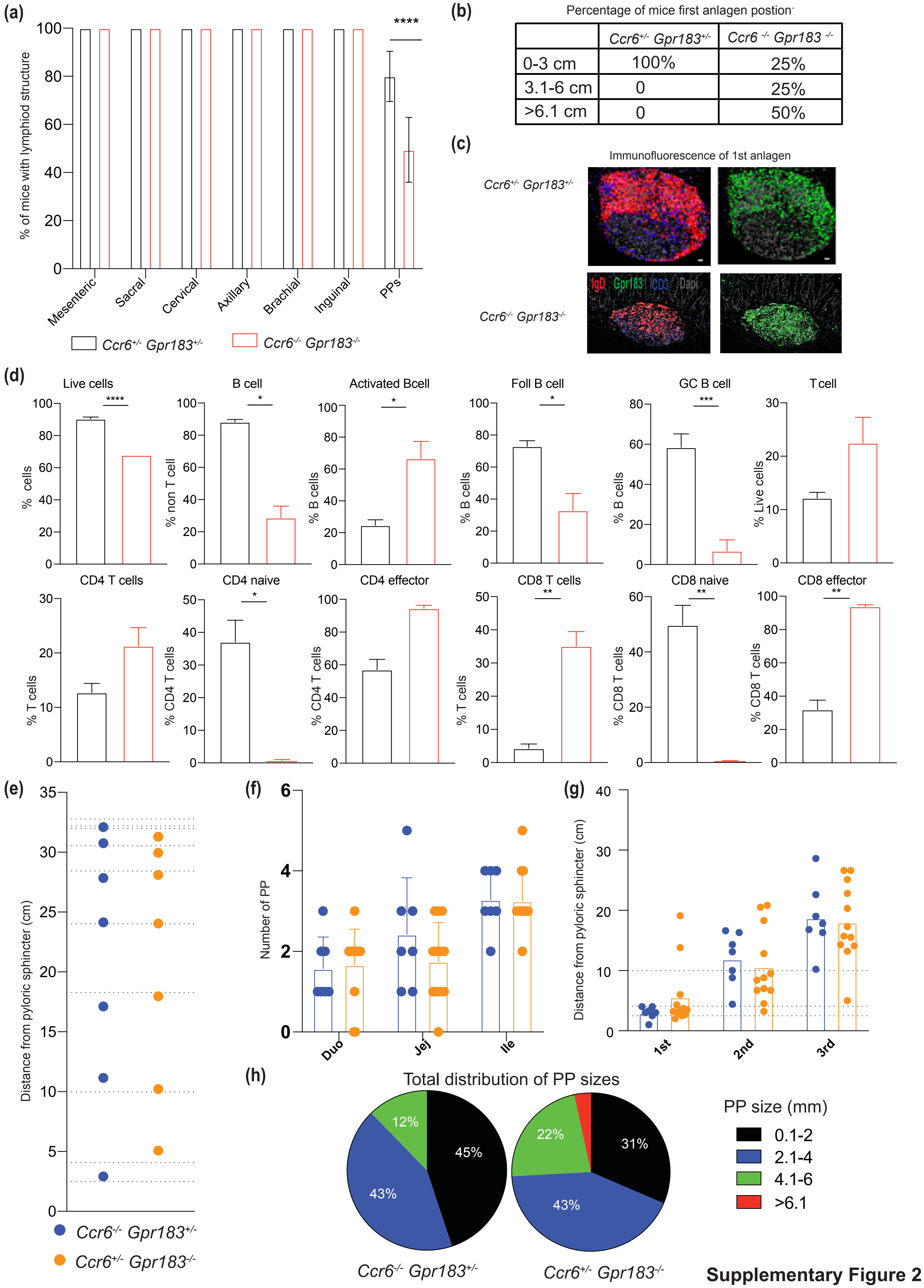


**Supplementary Figure 2**

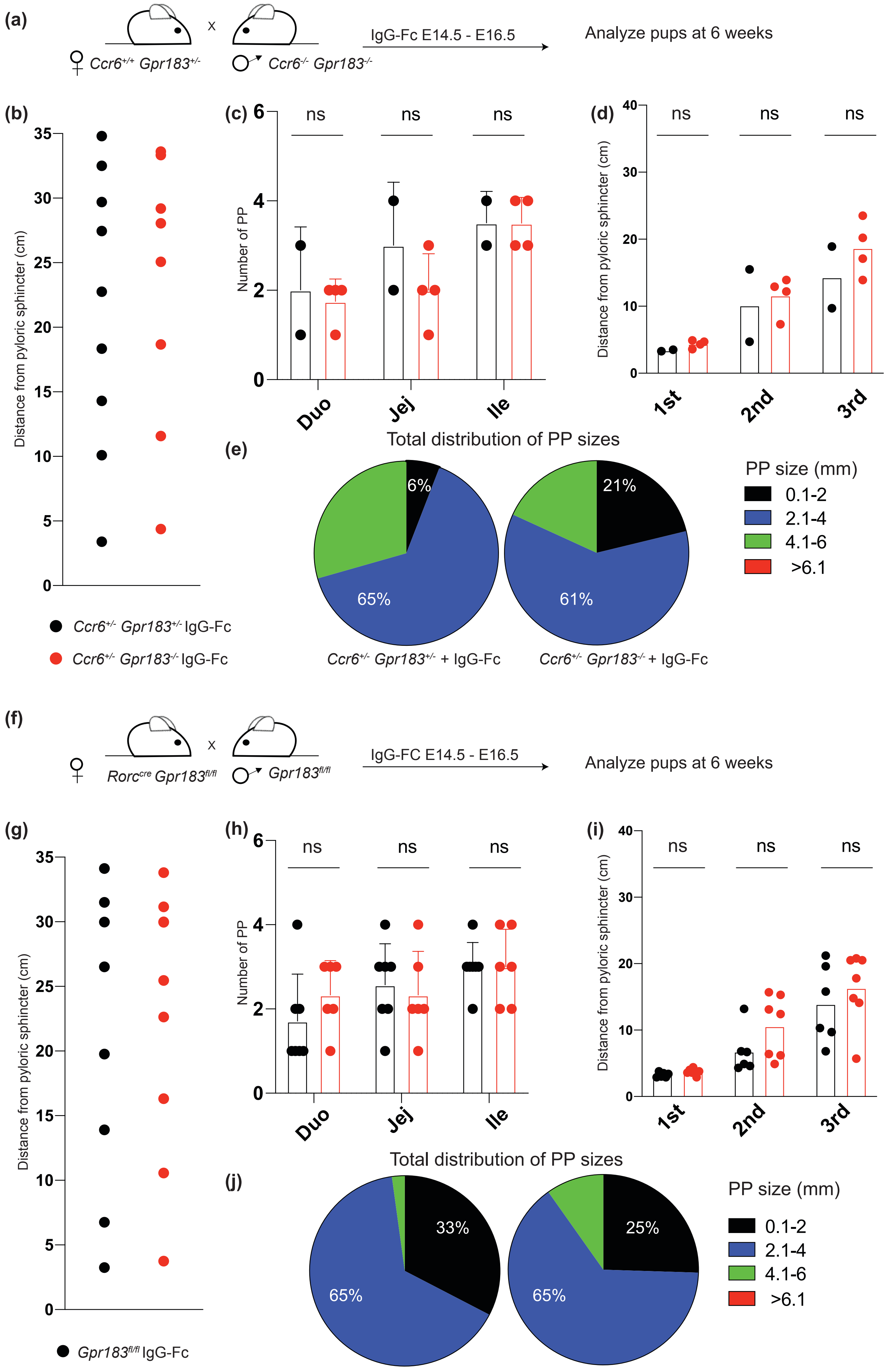


**Supplementary Figure 3**

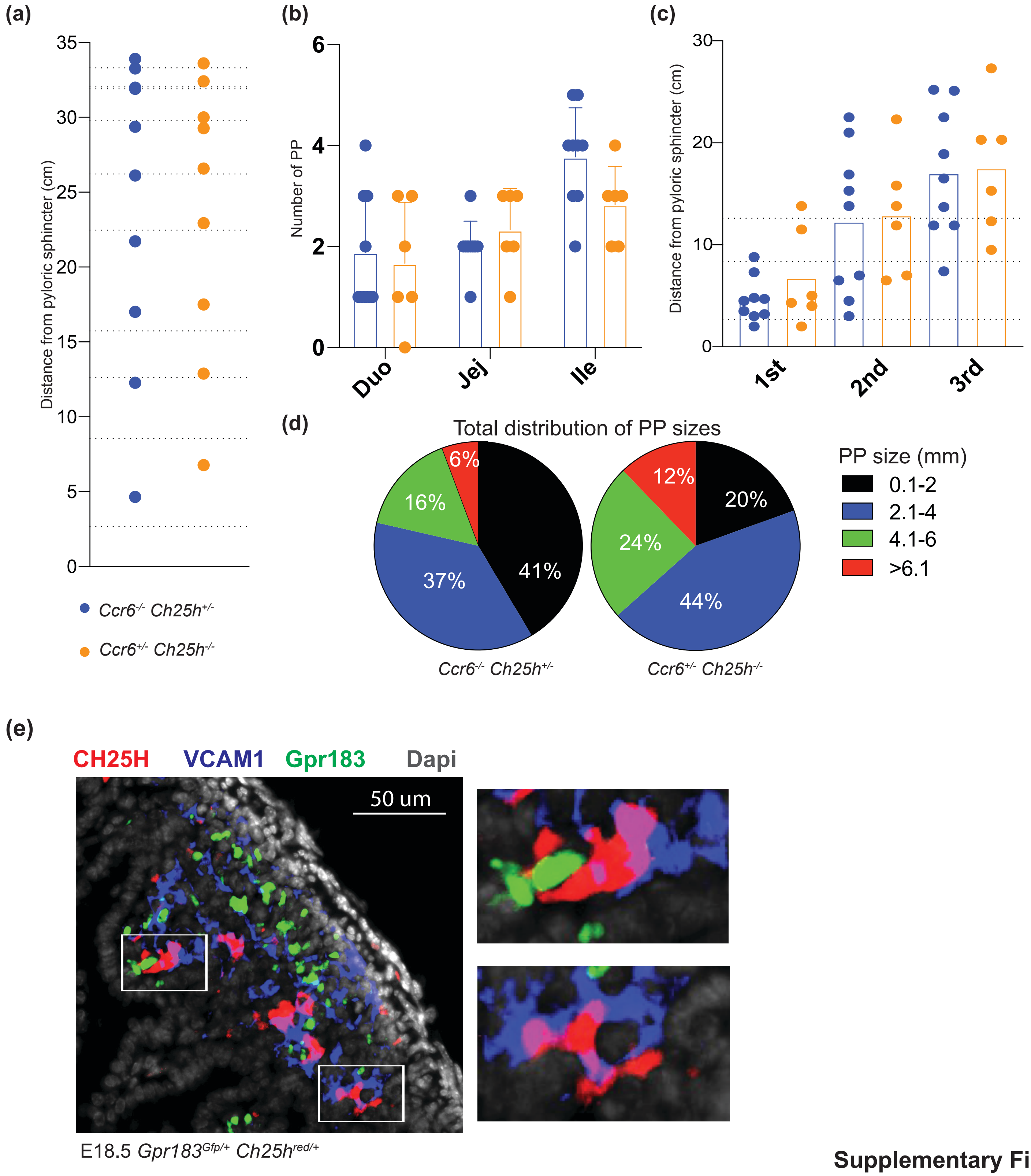


**Supplementary Figure 4**
